# Untangling the animacy organization of occipitotemporal cortex

**DOI:** 10.1101/2020.07.17.206896

**Authors:** J. Brendan Ritchie, Astrid A. Zeman, Joyce Bosmans, Shuo Sun, Kirsten Verhaegen, Hans P. Op de Beeck

**Affiliations:** Laboratory of Biological Psychology, Department of Brain and Cognition, Leuven Brain Institute, KU Leuven, 3000 Leuven, Flemish Brabant, Belgium; Facutly of Medicine and Health Sciences, University of Antwerp, 2000 Antwerp, Antwerp, Belgium

**Author notes:** Corresponding Author: J. Brendan Ritchie. Address: Psychologisch Instituut, Tiensestraat 102 – box 3714, 3000 Leuven, Belgium.

**Keywords:** category selectivity, animacy, faces, bodies, object categorization, fMRI, representational similarity analysis, occipitotemporal cortex, deep learning

## Abstract

Some of the most impressive functional specialization in the human brain is found in occipitotemporal cortex (OTC), where several areas exhibit selectivity for a small number of visual categories, such as faces and bodies, and spatially cluster based on stimulus animacy. Previous studies suggest this animacy organization reflects the representation of an intuitive taxonomic hierarchy, distinct from the presence of face- and body-selective areas in OTC. Using human fMRI, we investigated the independent contribution of these two factors – the face-body division and taxonomic hierarchy – in accounting for the animacy organization of OTC, and whether they might also be reflected in the architecture of several deep neural networks. We found that graded selectivity based on animal resemblance to human faces and bodies masquerades as an apparent animacy continuum, which suggests that taxonomy is not a separate factor underlying the organization of the ventral visual pathway.

## Introduction

One of the most fascinating examples of functional specialization in the human brain is the presence of areas in lateral and ventral occipitotemporal cortex (OTC) that preferentially respond to a small number of ecologically important visual categories including: faces (Kanwisher et al. 1997; McCarthy et al. 1997; Tsao et al. 2006), bodies (Downing et al. 2001; Bracci et al. 2010), places (Aguirre et al. 1998; Epstein and Kanwisher, 1998; Kornblith et al. 2013), tools (Martin et al. 1996), words (Baker et al. 2007; Cohen et al. 2000), and numerals (Shum et al. 2013). Beyond their importance to categorizing what we see, these areas have also been proposed as part of the neural substrate for the visually derived aspects of our conceptual or “semantic” knowledge of the world (Barsalou et al. 2003; Martin, 2016; Ralphs et al. 2017).

Of course, the category selectivity of these areas reflects only a small fraction of the things we know about so it is significant that OTC may represent more abstract properties of objects as well. At a broader spatial scale, the areas cluster in a manner that respects the superordinate dichotomy between animate objects that are capable of volitional self-movement and inanimate objects that are not (Bao et al. 2020; Behrmann and Plaut, 2013; Grill-Spector and Weiner, 2014). For example, face and body areas cluster separately from those for places and tools. However, some studies suggest OTC also represents animacy in a continuous, or even hierarchical, fashion (Sha et al. 2015; Thorat, Proklova and Peelen, 2019). In these studies stimuli consist of animal images grouped based on an intuitive taxonomy in which some animals rank high on the animacy scale (e.g. primates), some are intermediary (e.g. birds), and others (e.g. insects) are low (Connolly et al. 2012, 2016; Nastase et a. 2017; Sha et al. 2015). These results introduce the possibility that OTC represents conceptual relations between categories that cannot be so easily reduced to their diagnostic visual properties (Bracci, Ritchie, and Op de Beeck, 2017; cf. Fairhall and Carammaza, 2013).

The face-body division and conceptual taxonomy may both be factors that help explain animacy organization in OTC. These factors are not mutually exclusive, but they are not the same. Images of the face and (face-cropped) body of a person sit at the same (high) taxonomic level, yet are distinct, resulting in clearly dissociable neural responses. To date, studies of the animacy organization have failed to disentangle these two factors in several ways. First, studies providing evidence of taxonomic organization in OTC have used images of whole animal bodies and ignored the face-body division in general, and more specific issues such as the fact that we may be more accustomed to looking at the faces of some animals, and the bodies of others. Thus, these studies are unable to determine how the taxonomic organization is related to the face-body division. Second, these studies have focused on large swaths of OTC, and so are unable to determine whether the taxonomic organization is exhibited more narrowly in category-selective areas, like those for faces and bodies. Finally, these studies equate the idea of a continuous, graded organization in OTC with the representation of a taxonomic hierarchy. Therefore, they do not allow for the possibility that the animacy continuum in OTC is better explained by gradation in the representation of animal faces and bodies. More concretely, the apparent animacy continuum may simply reflect the relative similarity of animal faces and bodies to those of humans, for which face and body areas are known to be preferentially selective.

In light of these shortcomings, we performed an fMRI experiment aimed at disentangling these two factors and their contributions to explaining the animacy organization of OTC. First, we designed a stimulus set that allowed us to evaluate whether the taxonomic hierarchy explains the relationship between activity patterns in OTC when controlling for the face-body division, and vice versa. Second, we investigated the influence of both factors in explaining the relationship between activity patterns in more circumscribed, category-selective areas of OTC. Third, we investigated whether face and body morphology – in particular, the relative similarity of animal faces and bodies to those of humans – might better explain the relationship between activity patterns than taxonomy. Finally, we assessed whether both factors might be reflected in the patterns of activation weights of layers of multiple deep neural networks (DNNs), which some studies suggest also distinguish the animacy of objects (Bracci et al. 2019; Jozwik et al. 2017; Khaligh-Razavi and Kriegeskorte, 2014; Zeman et al. 2020).

## Results

We carried out an fMRI experiment in which participants (N = 15) viewed 54 natural images while in the scanner for two sessions. The stimuli included a face and body image of 24 animals, for a total of 48 animal images. The animals depicted were selected to form a taxonomic hierarchy following the stimulus design of Sha et al. (2015). The groupings included: two clusters of mammals based on intelligence (mammals 1 and mammals 2; see Materials and Methods), birds, reptiles/amphibians, fish, and invertebrates with exoskeletons (Figure 1A).

**Figure 1.**
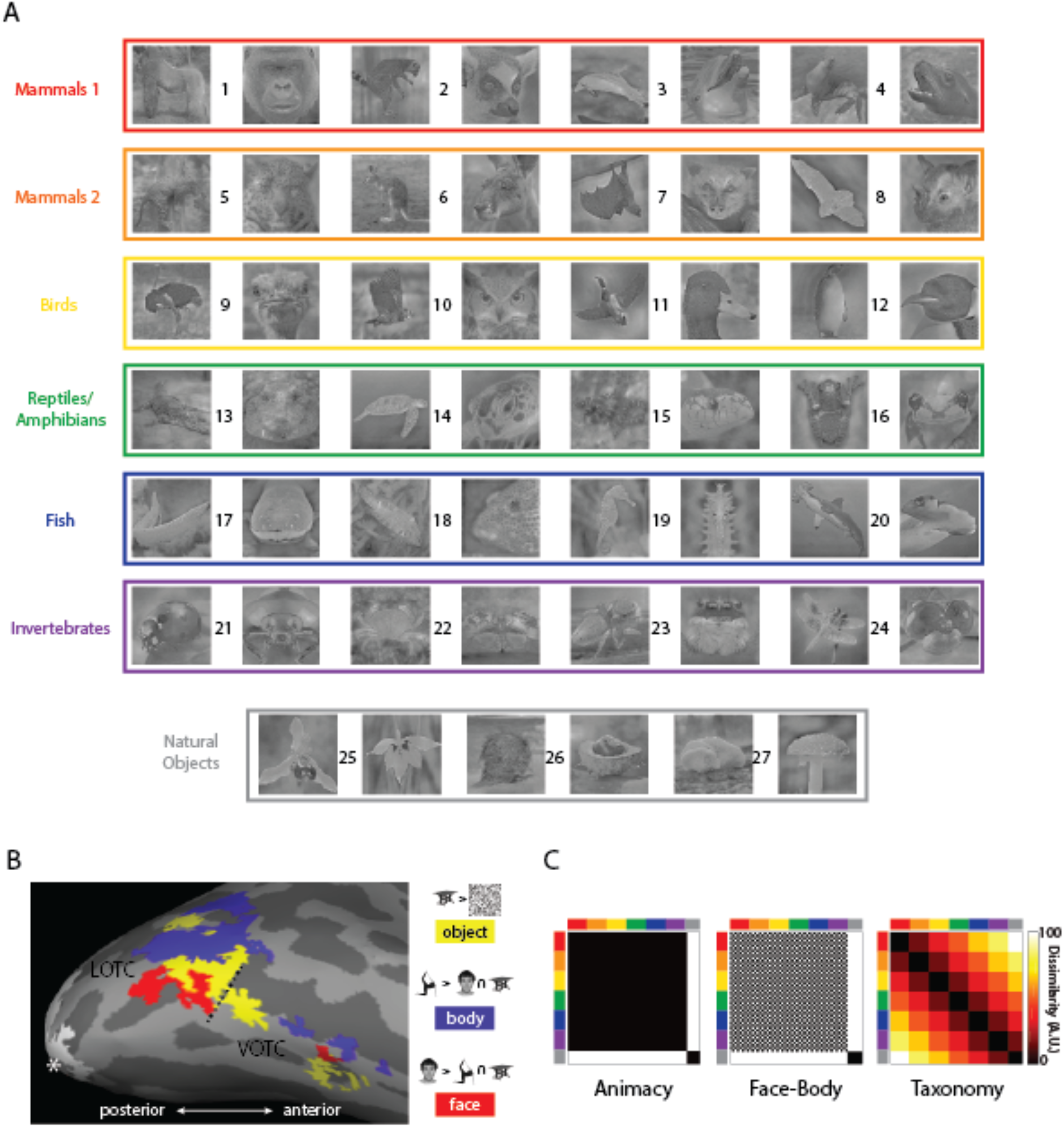
Core features of the experimental design. **(A)** All 54 natural image stimuli color-coded based on the taxonomic hierarchy. Numbers identify individual animals/natural object types: (1) gorilla, (2) lemur, (3) dolphin, (4) seal, (5) leopard, (6) kangaroo, (7) flying fox, (8) bat, (9) ostrich, (10) owl, (11) duck, (12) penguin, (13) crocodile, (14) turtle, (15) snake, (16) frog, (17) eel, (18) reef fish, (19) sea horse, (20) shark, (21) ladybug, (22) crab, (23) spider, (24) dragon fly, (25) orchid, (26) fruit/vegetable, and (27) mushroom. **(B)** The results of the functional contrasts used in the study to define the ROIs, for one representative participant, mapped onto their inflated cortex using FreeSurfer (Fischl, 2012). The hashed line indicates the boundary between the lateral and ventral masks defined using the Anatomical Toolbox (Eickhoff et al. 2005). The white asterisk indicates the occipital pole and the white patch is early visual cortex (EVC). **(C)** The three main model RDMs used throughout the study. The axes of the RDMs are color-coded to reflect the taxonomic hierarchy for the stimuli.

The remaining 6 images were of natural objects, with 2 images each of orchids, fruit/vegetables, and mushrooms. The primary regions of interest (ROI) were face-, body-, and object-selective patches of lateral OTC (LOTC) and ventral OTC (VOTC) based on contrasts from functional localizer scans (Figure 1B). We carried out representational similarity analysis (RSA) in order to compare the neural responses from these ROIs with models of the stimuli and behavioral judgments (Kriegeskorte and Kievet, 2013; Kriegeskorte, Mur, and Bandettini, 2008). RSA involves constructing representational dissimilarity matrices (RDM) from data of multiple modalities, in which each matrix cell reflects the pairwise dissimilarity between stimuli based on some metric. To disentangle the main factors related to animacy, model RDMs were constructed for the animate-inanimate division (Animacy), the face-body division (Face-Body), and the taxonomic continuum (Taxonomy), and compared with the neural RDMs for the different ROIs (Figure 1C).

### Face-body and taxonomy models both uniquely explain neural similarity in OTC

We first sought to determine whether the Animacy, Face-Body, and Taxonomy model RDMs would correlate with the neural RDMs for LOTC and VOTC, broadly construed. To this end we grouped all lateral and ventral ROIs in OTC (face, body, and object) into two broad ROIs “LOTC-all” and “VOTC-all”. The model RDMs were correlated with the neural RDMs of individual subjects for both OTC ROIs and early visual cortex (EVC), which was treated as a control region (Figure 2A). All statistical comparisons were made with a non-parametric Wilcoxon Signed Rank Test. All of the median correlations of the three model RDMs were significant for both LOTC-all and VOTC-all (all W(15) > 100, p < 0.01), but not EVC significant (all W(15) < 85, p > 0.15). For LOTC-all, there were also significant differences between the median correlations for Animacy vs Face-Body and Face-Body vs Taxonomy (both W(15) > 117, p < 0.001). Despite similar looking median correlations for VOTC-all, these differences were not significant (both W(15) < 67, p > 0.06).

**Figure 2.**
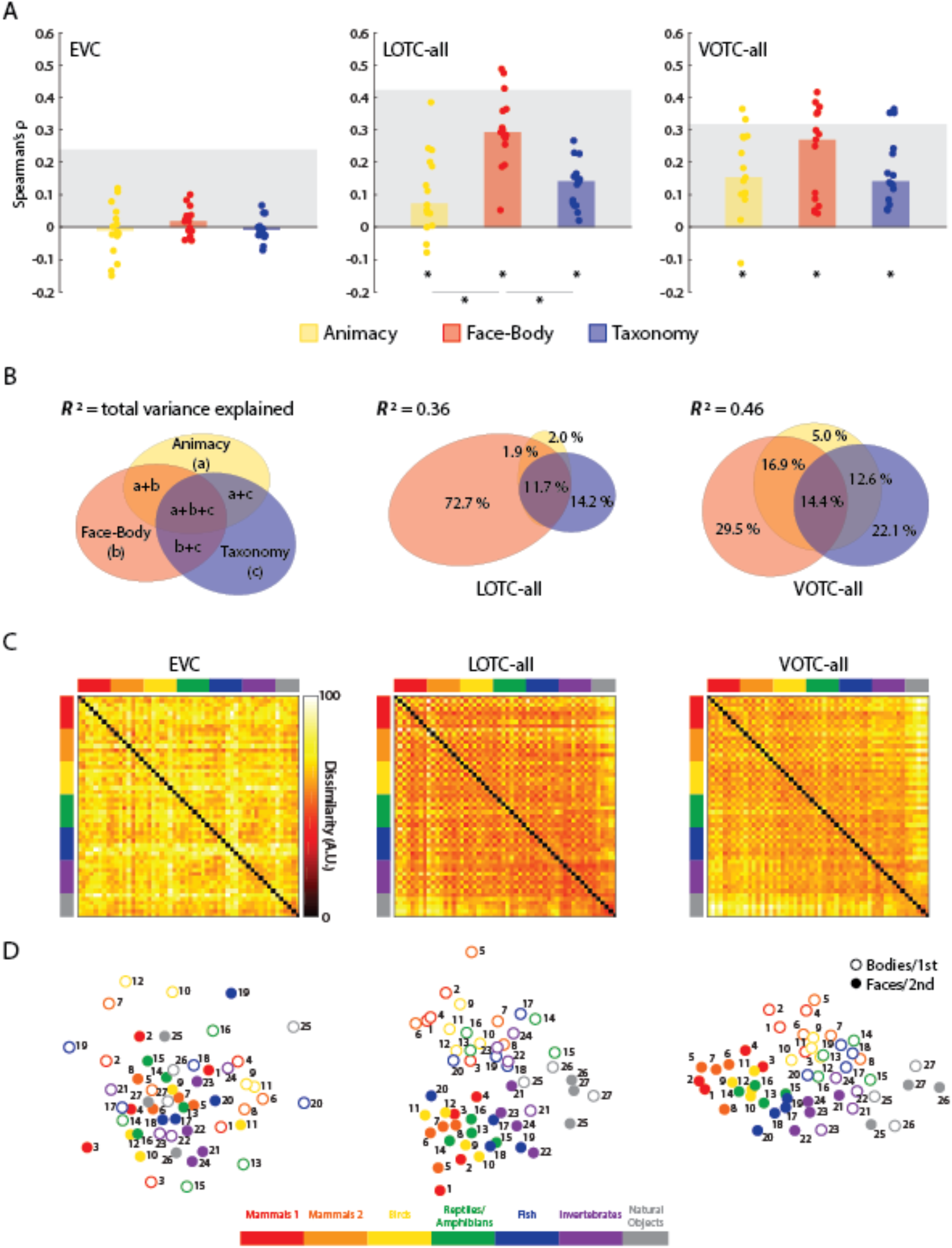
Comparing the main model RDMs to EVC, LOTC-all, and VOTC-all. **(A)** Bar charts indicating the median Spearman’s ρ rank-order correlations between neural RDMs for individual subjects (dots) and model RDMs. Gray blocks indicate the noise ceiling. * = p < 0.05. **(B)** Results of commonality analysis for LOTC-all and VOTC-all, depicted with Euler diagrams. A schema of the unique and common variance components is also depicted. Coefficients of determination (*R*^2^) indicate the total proportion of explained variance for the full model. Regions of the Euler plots indicate percentages of the explained, and not total, variance accounted for by each component. **(C)** Group-averaged neural RDMs with the axes color-coded based on the taxonomic hierarchy. Dissimilarity values are scaled to range 0-100. **(D)** 2D multi-dimensional scaling applied to the dissimilarity matrices. Points are color-coded to reflect the taxonomic hierarchy and are either rings or dots to reflect the face/body division, or the 1^st^/2^nd^ item for each natural object type. Numbers indicate individual animals or natural object type based on Figure 1A.

A number of nuisance models were also considered to determine whether they might also account for the structure of the neural RDMs for EVC, LOTC-all, and VOTC-all. First, it has been suggested that low- and mid-level image properties may account for apparent effects of object category in OTC (Andrews et al. 2015; Coggan et al. 2016; Long, Yu, and Konkle, 2018; Rice et al. 2014). To determine whether such properties might also account for the present results we used GIST, which captures spatial frequency and orientation information of images (Oliva and Torralba, 2001). Second, to ensure that familiarity was not a confound, prior to all experiments the subjects performed a naming task (see Materials and Methods). Finally, in order to maintain attention, during scanner subjects performed a one-back image preference task. RDMs for these tasks and GIST were at most weakly correlated the main model RDMs (Figure S1) and could not account for the results observed for OTC. There was a significant median correlation between GIST and the individual RDMs for EVC (Figure S2).

As suggested by Figure 1C, all three model RDMs are correlated with each other (Figure S1), which raises the issue of how much of the observed effects for the three models are unique to each of them or reflect their common structure. To address this, we carried out commonality analysis on the group-averaged neural RDMs for LOTC-all and VOTC-all. Commonality analysis is a method for determining how much of the explained variance of a linear model containing multiple predictors is unique to each predictor or is shared (Newton and Spurell, 1967; Siebold, McPhee, 1979) and can be used in conjunction with RSA (Groen et al. 2018; Hebart et al. 2018; Lescroart et al. 2015). Briefly, a vector of the coefficients of determination (*R*^2^) for all possible regression models (containing one, two, or three of the models as predictor) are differentially weighted to partition the explained variance of the “full” model (which contains all three models as predictors) into uniquely or commonly explained components (see Materials and Methods). Results are visualized using Euler plots (Figure 2B), in which areas of overlapping ellipses are proportional to the uniquely or commonly explained variance (Groen et al. 2018; Micallef and Rodgers, 2014).

The full model, containing all three predictors (Animacy, Face-Body, Taxonomy), explained a sizable amount of the variance for both ROIs (Figure 2B) and was significant based on permutation tests (LOTC-all: *R*^2^ = 0.36, p = 0; VOTC-all: *R*^2^ = 0.46, p = 0). Very little of this variance was uniquely explained by the Animacy model (Figure 2B). In LOTC-all, most of the explained variance was uniquely accounted for by the Face-Body model and to a much lesser extent by the Taxonomy model. Some variance was accounted for by all three, which likely reflects the shared structure between all three models described earlier. For VOTC-all, the Face-Body and Taxonomy were qualitatively more equitable in their unique contributions, with similar amounts of explained variance accounted for by the different common components. Crucially, these results show that the Face-Body and Taxonomy models each account for unique and independent components of the explained variance in neural dissimilarity in lateral and ventral OTC for our image set.

The interpretation of these results is further aided by visualizing the group averaged neural RDMs (Figure 2C) and comparing them to the model RDMs (Figure 1C). When this is done, it is clear that the RDMs for LOTC-all and VOTC-all, but not EVC, show structural similarity to the main model RDMs. Multidimensional scaling also makes more salient the differences in the commonality analysis results of LOTC-all and VOTC-all (Figure 2D). On the one hand, the face-body division is much more pronounced in the 2D space for LOTC-all. On the other hand, the taxonomic hierarchy is more apparent in VOTC-all, while the face-body division is also still clearly present.

### Face-body and taxonomy models both explain neural similarity in face- and body-selective areas of OTC

OTC is well-known to contain regions that show preferential selectivity for face and body images in both lateral and ventral OTC, as well as selectivity for objects more generally (Grill-Spector et al. 1999). Therefore, we sought to assess whether the pattern of results observed in LOTC-all and VOTC-all might be maintained in subordinate face- and body-selective areas. To this end we carried out the same analysis as before: correlating individual neural RDMs for LOTC-body and –face, and VOTC-body and –face areas with the three model RDMs, followed by commonality analysis. These analyses were then also carried out for both LOTC-object and VOTC-object areas.

For all three model RDMs (Figure 3A) the median correlations across subjects were significantly above chance for all four ROIs (all W(15) > 108, p < 0.01). For both LOTC-body and LOTC-face there were also significance differences between the median model correlations (all W(15) > 87, p < 0.02). In contrast in VOTC-body neither comparison was significant (both W(15) < 7, p > 0.44), and only Face-Body vs Taxonomy (W(15) = 88, p = 0.010), and not Animacy vs Face-Body (W(15) = 42, p = 0.25), showed a significant difference in median correlation in VOTC-face. We also found that somewhat similar results could be obtained when using the absolute difference in univariate responses as the dissimilarity metric (Figure S3). Still, while the face-selective areas showed substantive median correlations with the Face-body model, the median correlations for the Taxonomy model were greatly diminished across all ROIs.

**Figure 3.**
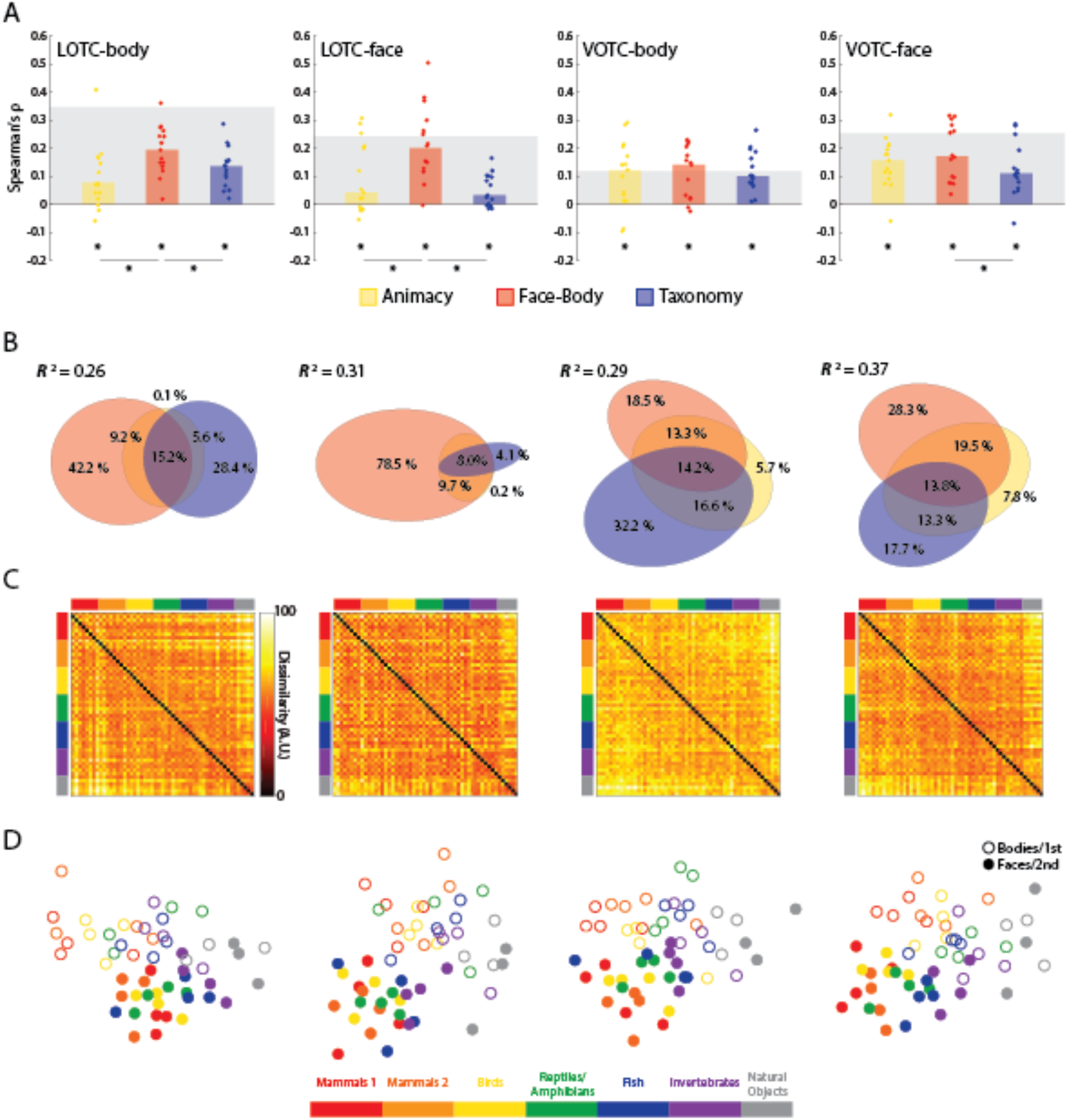
Results of model RDM comparisons to face and body-selective regions of OTC. **(A)** Bar charts indicating the median Spearman’s ρ rank-order correlations between neural RDMs for individual subjects and the three main model RDMs, for all four ROIs. **(B)** Results of commonality analysis for all ROIs, visualized with Euler diagrams **(C)** Group-averaged neural RDMs for the four ROIs. **(D)** 2D multi-dimensional scaling applied to the dissimilarity matrices for each ROI. Conventions follow those of Figure 2.

When all three predictors were regressed on the group-averaged neural dissimilarities (Figure 3B), the full model explained significant variance in the group neural RDMs (all *R*^2^ > 0.25, p = 0). There was a notable difference however in the portioning of the explained variance between the lateral and ventral ROIs (Figure 3B). In LOTC-body, while a large portion of the explained variance was due to the Face-Body model, a sizable portion was also uniquely explained by Taxonomy as well, with virtually no unique contribution by Animacy. In LOTC-face virtually all the explained variance was uniquely accounted for by Face-Body. For the ventral areas, both Face-Body and Taxonomy, and their common components, were substantive contributors to the explained variance. The most notable difference between the two areas was that, in VOTC-body, Taxonomy was the best unique and combined predictor, while in VOTC-face it was Face-Body.

These results comport with those for LOTC-all and VOTC-all: stronger effects of the Face-Body model in the lateral ROIs, while the effects of Face-Body and Taxonomy were more equitable in the ventral ones. Some of the characteristics of the group averaged RDMs for the face- and body-selective areas include the hot bands for the division between animal and natural objects and the face-body checkering (Figure 4C), and MDS plots again show a clear face-body division as well as a taxonomic continuum (Figure 4D). The same pattern of results (for median correlations, commonality analysis, and visualization), including the contrast between the lateral and ventral regions, was also observed for adjacent object-selective regions LOTC-object and VOTC-object (Figure S4).

**Figure 4.**
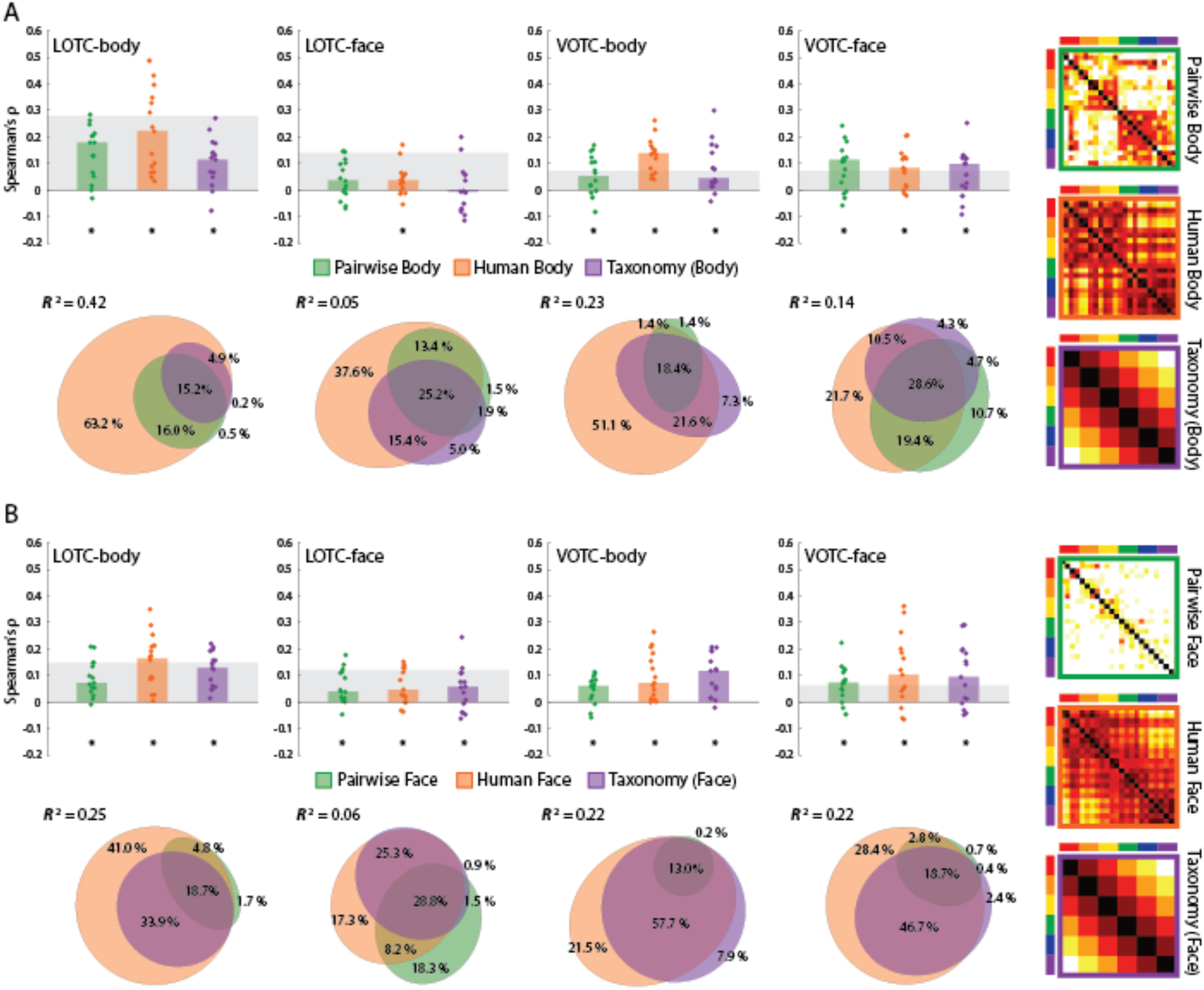
Results of comparing model RDMs for face and body stimuli to face and body-selective regions of OTC. **(A)** Bar charts indicating the median Spearman’s ρ rank-order correlations between neural RDMs for individual subjects and three model RDMs for the body images (also depicted): pairwise body similarity judgments (Pairwise Body), human similarity judgments (Human Body), and a (6-level) Taxonomy model for 24 images. **(B)** Bar charts and commonality analysis for the three model RDMs for the face images (also depicted). All conventions are the same as in Figures 2.

### Humanness, not taxonomy, best explains neural similarity in face- and body-selective areas of OTC

Next, we investigated the neural dissimilarity of face and body images separately in each of the face- and body-selective ROIs. For the images two kinds of similarity judgments were collected (Figure S5). First, separate groups of subjects judged the pairwise similarity of face or body images based on a 6-point scale (1 = high similarity). Second, other groups of subjects judged how similar the face or body images were to a human face or body. The inclusion of these human similarity tasks was inspired by the finding that relative similarity to humans may account for the animacy continuum (Contini et al. 2019), and that this may in fact reflect gradation in selectivity for animal faces and bodies. These judgments were converted to group averaged behavioral RDMs, which were compared to a Taxonomy model RDM constructed for just the 24 face and body images that included the six levels of the intuitive taxonomic hierarchy (Figure 1A).

The two behavioral RDMs for the body images and the 24-item Taxonomy model RDM were all highly correlated with each other (all ρ > 0.43, p < 0.0001). The two behavioral RDMs for the face images and truncated Taxonomy model RDM were likewise highly correlated (all ρ > 0.4, p < 0.0001). As seen in Figure 4A-B, the human body RDM suggests a grouping of the mammals and birds separate from the reptiles/amphibians, fish, and invertebrates, with a similar division for the human face RDM, though with greater similarity between the judgments for the bird and reptile/amphibian faces. These aspects that differentiate the human similarity matrices from the taxonomy model are reliable, given that our estimate of the split-half reliability of these matrices was r = .92 (human body) and r = .89 (human face), and so was much higher than their respective correlations with the Taxonomy model.

We correlated the pairwise similarity, human similarity, and Taxonomy RDMs with the separate neural RDMs for the face and body images across the face- and body-selective ROIs. For the body images (Figure 4A), there were significant median correlations between each of the model RDMs and individual neural RDMs for all but one ROI (all W(15) > 100, p < 0.02). The exception was LOTC-face where a significant median correlation was only observed for the human body RDM, but not the other matrices (both W(15) < 92, p > 0.08). For the face images (Figure 6B), all three predictor RDMs showed significant median correlations across all four ROIs (all W(15) > 103, p < 0.02).

To determine the unique vs common contributions of the three predictors, we carried out commonality analysis across image types and ROIs (Figure 6A and 6B). The full model explained a significant amount of the variance across ROIs for the body images (all 0.04 < *R*^2^ < 0.42, p < 0.01) and face images (all 0.05 < *R*^2^ < 0.26, p = 0) based on permutation tests. For the body images, across all ROIs, the human body similarity judgments were consistently the best unique predictor of variance with both pairwise similarity and Taxonomy predicting very little of the remaining variance uniquely. For the face images, the same picture emerged, with the exception of LOTC-face where pairwise face judgments uniquely predicted slightly more of the variance. Taken as a whole, these results suggest that any apparent effect of the Taxonomy model is almost entirely a reflection of commonly explained variance with the human face and body similarity judgments.

### Face-body and taxonomy, but not animacy, models explain activation dissimilarity in DNNs

DNNs have been a game changer in artificial intelligence based on their ability to match human performance on image classification tasks. Most network architectures consist of numerous convolutional layers, which tend to abstract local feature information when trained on large stimulus sets, followed by a few fully connected (FC) layers that more closely mirror the category structure in the training set. Several studies have reported correlations between FC layers and category-selective areas in human OTC suggesting that the animacy division is represented in these networks when trained on natural images (Bracci et al. 2019; Jozwik et al. 2017; Khaligh-Razavi and Kriegeskorte, 2014; Zeman et al. 2020). However, these studies did not include a taxonomic hierarchy in their stimulus designs. Therefore, we correlated the layer RDMs for several well-known DNNs to the main model RDMs, the broadly defined ROIs, and finally the similarity judgment RDMs for the face and body images. In particular, we included: CaffeNet, the Caffe implementation of AlexNet (Krizhevsky, Sutskever, and Hinton, 2012; Jia et al. 2014), VGG-16 (Simonyan and Zisserman, 2015), GoogLeNet (Szegerdy et al., 2015), and ResNet50 (He et al. 2015). We also included CORnet (Kubilius et al. 2018), a relatively shallow recurrent network that purports to exhibit the best matches to the representational structure of multiple neural datasets (Schrimpf et al. 2018).

Across all five networks, the correlations with the Face-Body model increased and peaked close to the first FC layers, or CORnet’s decoding layer, followed by the Taxonomy model correlations peaking at the final FC/Decoding layers (Figure 5A). Importantly, the networks had no knowledge about the taxonomic relations between the animals depicted in our stimulus images. Therefore, the fact that a significant correlation was observed with the Taxonomy model suggests that similar structure can be derived based purely on visual features of the images. In marked contrast to the results of previous studies, the layer RDMs tended to be negligibly, or even negatively, correlated with the Animacy model. The layers of the network were also correlated with individual neural RDMs for the three initial ROIs (Figure 5B). The peak median correlations with EVC tended to occur for middle conv layers, while those for LOTC-all and VOTC-all tended to occur at the later conv or FC/decoding layers. These findings are consistent with the claim that FC/decoding layers better reflect the structure of regions of OTC.

**Figure 5.**
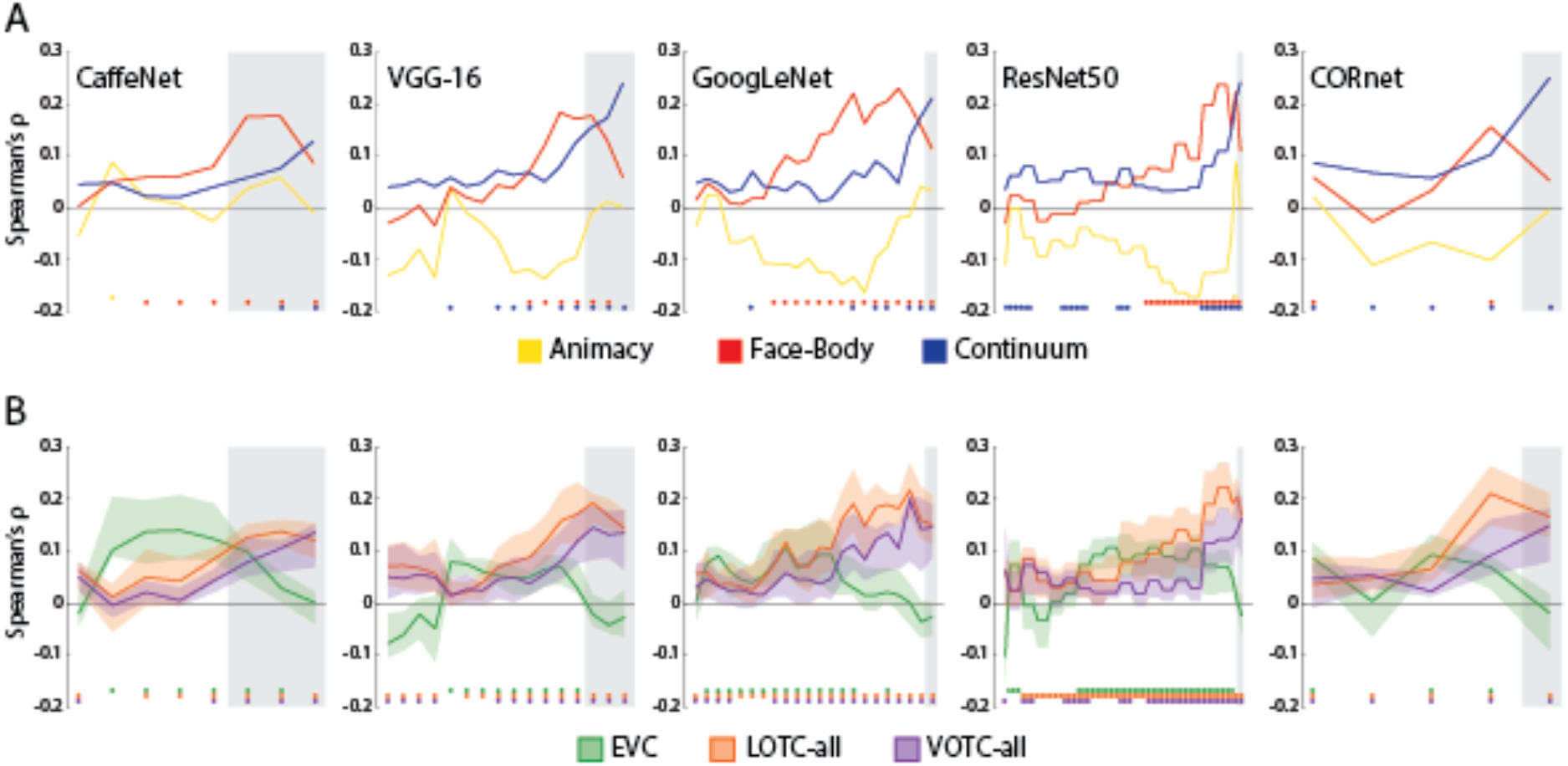
Results of RSA with DNNs for all 54 image. **(A)** the layers of each of the five networks correlated with each of the three model RDMs **(B)** The layers of each of the five networks correlated with the individual subject neural RDMs for the three initial ROIs considered. The solid lines indicate the median correlations at each layer, and the transparent regions range from the 1^st^ quartile to the 3^rd^ quartile. For both **(A)** and **(B)** Grey areas indicate FC/decoding layers for each network.

**Figure 6.**
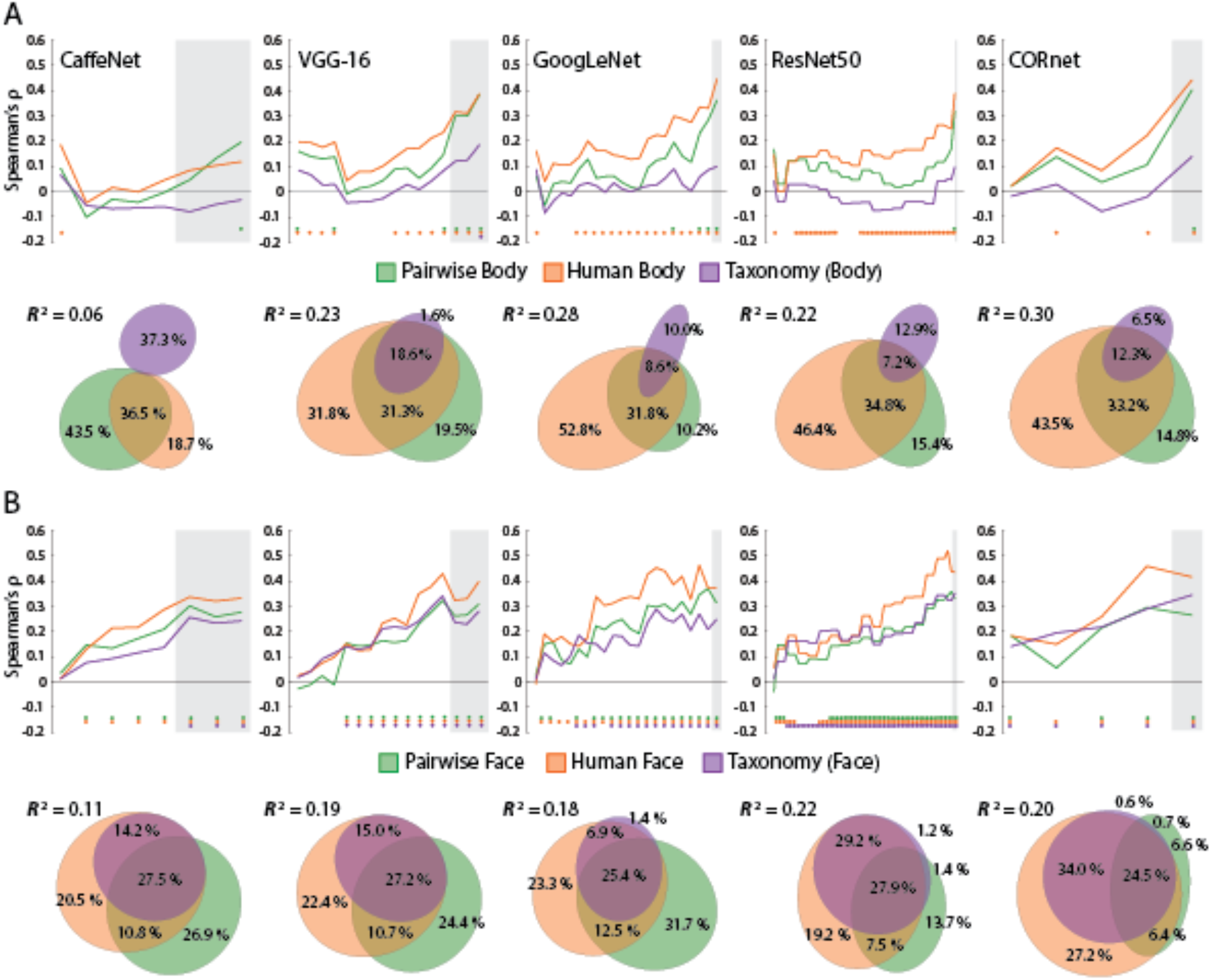
Results of comparing models for face and body images to DNNs. **(A)** the layers of each of the five networks were correlated with each of the body image RDMs. Color-coded dots indicate significant (positive) correlations at p < .05 (FDR-adjusted), based on two-sided permutation tests. Results of commonality analysis for the final layers of all networks are visualized with Euler diagrams. **(B)** Correlations between the layers of the five networks and the three model RDMs for the face images. All conventions are the same as in **(A).**

We also investigated whether the dissimilarity structure of the network layers might be better captured by pairwise and human similarity judgment RDMs for the face and body images. For the body image models (Figure 6A), we found that for all but one network (CaffeNet) the human body RDM tended to show the highest correlations across layers of networks, with the pairwise body RDM peaking at a similar, or higher level, at the final layers. In contrast, the Taxonomy model only showed a significant correlation with the final layer of VGG-16. For the face image models (Figure 6B), all three models showed a consistent increase in correlation effect sizes with network depth. Again, as with the neural data, the human face model tended to show the highest correlations. The Taxonomy model also consistently correlated with many of the layers of the different networks. These results suggest that correlation with the Taxonomy model for the full stimulus set (Figure 5A) was likely being driven by the face images.

To determine the unique vs common contributions of the three predictors we carried out commonality analysis across image types and the final layers of the DNNs (Figure 6A and 6B). We focused on the final layers of the networks because these consistently showed the peak correlation with the Taxonomy model for the full image set (Figure 5A). For the body images, the full model explained a significant amount of the variance for the final layers of all networks (0. 05 < *R*^2^ < 0.3, all p < 0.004). For all but one network, the human body RDM was consistently the best unique predictor, followed by the pairwise body RDM. The exception was CaffeNet, where comparatively little of the variance was explained by the full model and therefore the results of the commonality analysis are difficult to interpret. For the face images, the full model explained a significant amount of the variance for the final layers of all networks (0.1 < *R*^2^ < 0.23, all p = 0). For ResNet50 and CORnet the human body similarity judgments were the best unique predictor of variance. For the remaining networks the pairwise face RDM explained the most unique portion of the variance of the full models. In turn, even though the Taxonomy model was significantly correlated with the final layer of the networks, it explained virtually no unique variance in the final layer dissimilarity values.

These findings are broadly consistent with those observed for the face- and body-selective areas (Figure 4). There are two notable differences: first, the pairwise similarity RDMs consistently rivaled, or surpassed, the human similarity RDMs as predictors of layer dissimilarity values; and second, the taxonomy model was only a significant predictor for layer dissimilarity values for the face images. Crucially, when it came to the final network layers, any effect of the Taxonomy model was enveloped by the variance components shared with the human face RDM. In sum, the apparent effect of Taxonomy in these networks is likely explained by the similarity of visual features between the face and body images, as captured by the pairwise and human similarity judgments.

## Discussion

Animacy has been proposed as an important organizing principle in OTC (Behrmann and Plaut, 2013; Grill-Spector and Weiner, 2014). Two factors influencing this organization are suggested by the literature: the face-body division and intuitive taxonomy. In the present study we sought to isolate the influence of these two factors and found that they independently explained the variance dissimilarity values in OTC. When OTC was partitioned, the same pattern of positive results was also observed in the face-, body-, and even object-selective areas. Crucially, human similarity judgments were however shown to be much better predictors than the Taxonomic model when the data for face and body images were analyzed separately. Finally, the later layers of DNNs also correlated with both the Face-Body and Taxonomy models – but not Animacy – and sensitivity to Taxonomy found in the final layers of the networks was better captured by the pairwise and human similarity judgments for the face and body images. These results have important implications for: (i) the supposed taxonomic organization of OTC; (ii) whether OTC in fact represents the animacy of objects; (iii) whether DNNs distinguish the animacy of objects in images; and (iv) the scope of face- and body-selectivity in OTC.

### Occipitotemporal cortex does not represent taxonomy

Previous studies suggest that animacy organization of OTC may reflect the representation of a continuum, rather than a dichotomy, between animate and inanimate objects. For example, Sha et al. (2015) found that neural dissimilarity in LOTC showed no such division. The stimuli used in their study were selected to form an intuitive taxonomic hierarchy with primates (human and chimpanzee) as the most animate compared to invertebrates (ladybug and lobster), with other mammals, birds, and fish in between. Other studies have also found that pattern similarity in OTC to reflect an ordering based on intuitive taxonomy. For example, Connolly et al. (2012) found that the taxonomic ordering between primates, birds, and insects was reflected in the area, and Connolly et al. (2016) found similar results with mammals, reptiles, and invertebrates. More recently, a study by Thorat et al. (2019) reported that neural dissimilarity in VOTC was well-captured by judgments of the relative capacity for thoughts and feelings, which they term “agency” — though notably these are properties of entity subjectivity not agency (Gray, Gray, and Wegner, 2007). Still, the resulting ranking of the images was very similar to what one would predict based on an intuitive taxonomic grouping of their object images.

Our results differ from these previous studies in a number of ways. First, we consistently found a robust animate-inanimate dichotomy in neural dissimilarity across ROIs, in direct contrast to the results of Sha et al. (who used tools as their contrast class). Second, none of these previous studies controlled for the face-body division in their stimulus designs. Therefore, it is significant that the Taxonomy model independently contributes to explaining the neural dissimilarity in OTC for both face and body images. This finding on its own can be interpreted as providing positive evidence of a taxonomic continuum in OTC. However, third, we also considered the possibility that the apparent continuum does not reflect representation in OTC of taxonomic relations per se, but rather graded responses to the images in face- and body-selective areas of OTC. To evaluate this alternative, we compared the neural dissimilarity of face- and body-selective ROIs to pairwise and human similarity judgments for the face and body images. Commonality analysis revealed that human similarity was the dominant predictor, accounting for sizable portions of uniquely explained variance for both face and body images across face- and body-selective regions.

Therefore, taken as a whole, our results suggest that the apparent animacy continuum does not reflect the representation of an intuitive taxonomy, or a correlated agency dimension. Instead, they support an alternative hypothesis: there is no taxonomic organization in OTC. Instead, as OTC is well-known to display distributed and differential selectivity for faces and bodies, the apparent continuum effect simply reflects variation in response of animal faces and bodies based on how similar they are to human faces and bodies, which are plausibly the most preferential stimuli of these categories for the ROIs. Of course, what accounts for the morphological differences between faces and bodies between species is their evolutionary history, and our intuitive grouping of animals based on their visual features in part reflects this. Still, the most parsimonious explanation of the available data is that OTC simply represents animal faces and bodies and how much they resemble a human face/body and not a taxonomic continuum per se.

### Occipitotemporal cortex does not represent object animacy

Given that the apparent animacy continuum may simply reflect gradation in the face- and body-selectivity throughout OTC, our results also raise the question of whether OTC represents animacy at all. Many of the studies on the animate-inanimate division have tended to focus on large swaths of OTC in spite of the well-known functional division between different category-selective areas. Yet, these areas themselves are spatially pooled depending on whether they represent animate stimuli. The clearest division in this respect is between face- and body-selective areas that are more lateral in the VOTC, compared to the more medial portions selective for scenes. Based on this, one possibility is that animacy is represented at a broader spatial scale, which subsumes areas that represent more specific animate or inanimate categories such as faces or scenes (Bao et al. 2020; Grill-Spector and Weiner, 2014). An alternative hypothesis is that the higher-level category of animacy is not represented at all, even though the topographic organization of category-selective areas reflects the animate-inanimate division. In other words, any apparent broad representation of animacy is an artefact of (principally) strong selectivity for faces and bodies. This face-body hypothesis seems to be supported by our finding that neural responses are explained better by human similarity than by taxonomy.

This somewhat deflationary hypothesis is also supported by the recent study of Bracci et al. (2019), who selected three groups of object images: animals, artefacts, and “look-alike” artefacts. These images were chosen to form trios, such as an image of a duck, a kettle, and a duck-shaped kettle. The stimulus design of Bracci et al. allowed for two possibilities: that neural dissimilarity would exhibit a division based on object appearance (animals and look-alikes vs artefacts) or object identity (animals vs look-alikes and artefacts). They found that neural dissimilarity in VOTC correlated with the object appearance, and not the object identity model. Based on this finding, Bracci et al. proposed that VOTC does not represent the animate-inanimate division, but selectivity for diagnostically important visual features. We would suggest that these features are specifically diagnostic for faces and bodies, as part of a feature-based neural code for object categories (Bracci, Ritchie, and Op de Beeck, 2017). One prediction of this deflationary view is that animals without faces or heads, or morphologically very different bodies, may not be differentiated in OTC from natural objects. Indeed, this possibility is already implied by the finding that, when subjects classify novel or unfamiliar objects as animals, plants, or minerals, they rely on midlevel visual features that tend to be diagnostic of animal body parts, such as symmetry (Schmidt, Hegele, and Fleming, 2017).

### Deep neural networks do not represent object animacy

DNNs are increasingly being used as models of visual processing (Cichy and Kaiser, 2019; Kriegeskorte, 2015; Serre, 2019). The use of DNNs in this capacity has in part been motivated by similarities in the activity patterns of the later FC layers to regions of OTC for networks trained on the ImageNet dataset. In particular, several studies using ImageNet trained networks have reported that FC layers exhibit a similar animacy organization to OTC, with a clear division between patterns of activation weights for images of animate and inanimate objects (Bracci et al. 2019; Jozwik et al. 2017; Khaligh-Razavi and Kriegeskorte, 2014; Thorat et al. 2019; Zeman et al. 2020).

Given these previous findings, it is striking that we did not observe a consistent animacy organization across five ImageNet-trained DNNs when they were presented with our image stimulus set. At the same time, our result should be less surprising when one considers that the Animacy model explained little of the unique variance in our OTC ROIs. The layers of the DNNs also did consistently correlate with the Face-Body and Taxonomy model RDMs. Given that there are morphological differences between the faces and bodies of different animals, it is likely that the networks are sensitive to these differences. Indeed, when the data for the face and body images were analyzed separately, we found that the pairwise and human similarity RDM explained most of the variance in the final layer RDMs. So, like with OTC, the ImageNet-trained DNNs plausibly do not represent either a taxonomic continuum or a categorical division between animate and inanimate objects, but rather graded representations of face- and body-related visual features. This is also consistent with the training history of these networks with still images, which provide information about how statistical properties of images relate to the categorization of the objects in these images yet contain no direct information about animacy. These findings are also more in line with the finding that, across many different networks, DNNs do not capture the structure of OTC when even simple manipulations are carried out on the images (Xu and Vaziri-Pashkam, 2020).

At the same time, we may predict different results when a network is explicitly trained on, for example, face stimuli, which could potentially help to promote the face-body distinction. Indeed, Hill et al. (2019) found that appropriately trained DNNs generated a multidimensional “face space” much like with human perception and found that the network could perform face identification on trained faces even when presented with caricatures. Thus, while the evidence that they represent animacy may in fact be overstated, DNNs may yet be shown to represent the same kinds of categories found in OTC, when appropriately trained.

### Rethinking face- and body-selectivity in OTC

Our results also have implications for how one conceives of the scope of face- and body-selectivity in OTC. Typically studies of face and body representation focus on human faces and bodies, with a notable exception being studies in which faces of conspecifics are included as stimuli. Some studies of the animacy organization in human and monkey OTC have included face and body images from multiple species, though without taking account of taxonomic relations (Cichy et al. 2014; Kiani et al. 2007; Kriegeskorte et al. 2008). The tacit methodological assumption is that human faces and bodies dominate the representational capacities of face- and body-selective areas in OTC, perhaps due to templates based on our own face and body morphology (de Haas, Schwarzkopf, Rees, 2016; Henriksson, Mur, and Kriegeskorte, 2015; Orlov, Makin, and Zohary, 2010; van den Hurk et al. 2015). In which case, we do not need to sample from the animal kingdom more broadly when trying to understand face or body-related topographic organization in OTC, or the structure of a “face space” (O’Toole et al. 2018). Yet, this simplifying assumption is surely mistaken, as we can readily recognize faces and bodies of other animals, including their morphological differences. For example, face-selective areas may encode a face template, but it will be influenced by our development during which we are typically exposed to many animals of all shapes, sizes, and facial features (Arcaro et al. 2017; Sugita et al. 2008).

The hypothesis that face- and body-selective areas are organized to also represent non-human faces and bodies provides one plausible explanation of why we so readily experience phenomena such as face pareidolia, in which inanimate objects look like they have faces. VOTC-face shows a preferential and differential response to pareidolic stimuli when matched for object type (Wardle, Seymour, and Taubert 2017), and behavioral evidence shows that monkeys also experience face pareidolia (Taubert et al. 2017). But the question remains as to why the phenomenon occurs. Part of the explanation may be a system tuned to avoid “misses” of face detection at the cost of more “false alarms”. However, if OTC is specialized to represent faces and bodies very broadly, and including animals with morphological characteristics quite dissimilar from those of humans, then phenomena like pareidolia may be a natural by-product. For example, a smiley face has more visual features in common with the face of an electric eel (Figure 1A), than the face of a human.

### Summary and conclusion

Animacy is claimed to be one of the organizing principles of OTC and may reflect either the representation of a taxonomic hierarchy, or underlying selectivity for faces and bodies. In the present study we attempted to disentangle these factors. Our results suggest that a graded selectivity for faces and bodies, in terms of how much they resemble a human face or body, may masquerade as an animacy continuum, suggesting that taxonomy is not a separate factor underlying the organization of OTC. Indeed, our results provide further support for the idea that OTC is not representing animacy per se, but simply faces and bodies as separate from other ecologically important categories of objects. In this respect our results provide new insights into the functional organization of the ventral visual pathway more generally.

## Materials and Methods

### Participants

The fMRI experiment included 15 adult volunteers (10 women; mean age = 24.2 years; age range 21 to 33 years). This number of participants is sufficient to have a power above 0.95 with reliable data that guarantee an effect size of d = 1. Based upon previous studies with similar amounts of data per stimulus and subject (i.e. 2 long scan sessions per participant; see Bracci & Op de Beeck, 2016, Bracci et al., 2019), we know that the distinction between animals and regular objects in representational similarity analyses has a very high effect size (typically with a Cohen’s *d* of 1 to 4 even in smaller ROIs). We also assessed the between-subject reliability of the neural data by calculating the noise ceiling for correlations with the RDMs of each ROI (see below).

A total of 42 volunteers participated in the different similarity judgment tasks (33 Women; mean age = 21.5; age range 18 – 35 years), with subjects randomly selected to participate in the pairwise body, pairwise face, human body, and human face similarity tasks. All volunteers were predominantly right-handed, had normal or corrected vision, and provided written informed consent for participation in the experiments. All experiments were approved by the ethics committee of UZ/KU Leuven and all methods were performed in accordance with the relevant guidelines and regulations.

### Stimuli

Stimuli consisted of 54 natural images of objects (Figure 1A) and included a body and face image of 24 animals, as well as two images each of three natural objects, resulting in 48 animal and 6 natural object images. The animal images were selected to cluster into six levels of an intuitive taxonomic hierarchy: mammal cluster 1, mammal cluster 2, birds, reptiles/amphibians, fish, and exoskeletal invertebrates. The distinction between the two mammal clusters was intended to distinguish high intelligence mammals, including primates (gorilla and lemur) and trainable aquatically mobile mammals (dolphin and seal), from comparatively less intelligent mammals that are terrestrial (leopard and kangaroo) or capable of flight (flying fox and bat). Natural object images were of orchids, fruit/vegetables, and mushrooms. All images were cropped to 700 × 700 pixels (subtending ~10 degrees of visual angle in the scanner), converted to grey scale, focus blurred in the background regions, and then filtered using the SHINE toolbox (Willenbockel et al. 2010) to equate the luminance histogram and the average energy at each spatial frequency. For the fMRI experiment, stimulus presentation and control were via a PC computer running the Psychophysical Toolbox package (Brainard, 1997), along with custom code, in Matlab (The Mathworks, Inc). For the similarity experiments, and the naming task (described below), presentation and control were via PC computers running PsychoPy2 (Peirce, 2007).

### Similarity Judgment Experiments

Participants were tasked with making similarity judgments based on the sequential presentation of pairs of either face or body stimulus images (Figure S5). For the pairwise face and pairwise body similarity tasks subjects responded using a 6-point scale (1 = highly similar, 6 = highly dissimilar) as to how similar the two animal faces/bodies were to each other. For the human face and body similarity tasks the subjects responded whether they considered the first (press 1) or second (press 2) animal face/body was more similar to the face/body of a human. Across all similarity tasks, the trial structure was virtually identical: the fixation cross appeared (1000 ms), then the first image (1000 ms), an inter-stimulus interval (1000 ms), and then the second stimulus (1000 ms). After the second stimulus disappeared from the screen, text appeared reminding participants of either the 6-point scale or the pairwise choice options. The next trial did not start until subjects made a response. All possible sequential pairwise combinations of images were presented during the experiment, in random order, with five rest breaks evenly spaced throughout.

### Naming Task

Prior to both the similarity and fMRI experiments, all participants carried out a naming task (Figure S5): a fixation cross appeared for 500 ms after which an image appeared for 1000 ms, then subjects typed, in English or Dutch, the name of the animal or natural object depicted in the image (e.g. “duck”/“eend”). If a participant recognized the animal/natural object, but could not remember the name, they were instructed to type “y”, and if they did not recognize it at all, to respond with “n”. After typing their response subjects pressed “enter” and the image and the correct English and Dutch labels appeared for 3000 ms before the next trial began. Correct responses were coded based on pre-determined labels, with any discrepancy marked as incorrect. For example, if a subject correctly labeled the body image for animal 24 as “dragonfly”, but the corresponding face image as simply “fly”, the latter response would be coded as incorrect. Only the naming data from fMRI participants was coded and analyzed for comparison with their neural data.

### Scanning procedures

The fMRI experiment consisted of two sessions of eight experimental runs followed by two localizer runs, for a total of 16 experimental and four localizer runs per subject, with one or two anatomical scans also collected for each participant. Using a rapid event-related design, each experimental run consisted of a random sequence of trials including two repeats of each of the 54 images and 18 fixation trials, for a total of 144 trials per run. Each stimulus trial began with the stimulus being presented for 1500 ms, followed by 1500 ms of the fixation bullseye. Subjects performed a one-back task, where on each trial they indicated with a button press whether they preferred looking at the current image or the previous one. Experimental runs had a total duration of 7 m 30 s. A fixation bullseye was centrally presented continuously throughout each run.

For the localizer runs a block design was used with four stimulus types: bodies, faces, objects, and box-scrambled versions of the object images, with 18 images of each stimulus type. Each image in a block appeared for 400 ms followed by 400 ms fixation with four repeats of each stimulus block type per run. All four series of image types were presented sequentially in each stimulus block in pseudorandom order, followed by a 12 s fixation block. Localizer runs had a duration of 8 m 0 s. in order to maintain their attention during a run, participants indicated when one of the images was repeated later in an image series, which occurred once each for two different randomly selected images per image type, per block. A fixation bullseye was centrally presented continuously throughout each run. For one subject the data for 3 of 4 localizer runs was used, since the data file for the remaining run was corrupted and unusable.

### Acquisition parameters

Data acquisition was carried out using a 3T Philips scanner, with 32-channel coil, at the Department of Radiology of the KU Leuven university hospitals. Functional MRI volumes were acquired using a 2D multiband T2*-weighted echo planar (EPI) sequence: MB = 2; TR = 2000 ms; TE = 30 ms; FA = 90 deg; FoV = 216; voxel size = 2 × 2 × 2 mm; matrix size = 108 × 108. Each volume consisted of 46 axial slices (0.2 mm gap) aligned to encompass as much of the cortex as possible and all of the occipital and temporal lobes. Typically, this resulted in the exclusion of the most superior portions of the parietal and frontal lobes from the volume. The T1-weighted anatomical volumes were acquired for each subject using an MP-RAGE sequence, 1 × 1 × 1 mm resolution.

### fMRI preprocessing and analysis

Preprocessing and analysis of the MRI data was carried out with SPM12 (v. 6906) using default settings unless otherwise noted. For each participant, functional fMRI volumes was combined from the two sessions (while preserving run-order) and slice-time corrected (indexing based on slice acquisition time relative to 0 ms, not slice order), motion corrected using the realign operation (using 4^th^ degree spine interpolation), and coregistered to the individual anatomical scan. For all these steps the transformations were estimated and saved to the image header files, before a single reslicing was carried out using the coregistration (reslice) operation. Functional volumes were then normalized to standard MNI space by first aligning the SPM tissue probability map to the individual subject anatomical scan, and then applying the inverted warp to the functional volumes. Finally, all functional volumes were smoothed using a Gaussian kernel, 4 mm FWHM.

After preprocessing, the BOLD signal for each stimulus, at each voxel, was modeled separately for the experimental and localizer runs using GLMs. For the experimental runs, the predictors for the GLM consisted of the 54 stimulus conditions and six motion correction parameters (translation and rotation along the x, y, and z axes). The time course for each predictor was characterized as two boxcar functions at the two stimulus onsets (duration = 1500 ms) convolved with the canonical hemodynamic response function. The GLM analysis produced one parameter estimate for each voxel, for each stimulus predictor, for each run. For the localizer runs the same modeling procedure was carried out for the 4 localizer conditions, with a single onset at the beginning of each image series (duration = 16 s) for each image type, for each of the four stimulus blocks.

### Defining ROIs

Three contrasts were used to specify functional ROIs based on the four conditions from the localizer runs (Figure 1B): body > face + objects; face > body + object; and object > scrambled. We used conjunctions of masks from the Anatomy Toolbox to isolate lateral (conjunction of bilateral hOc4lp, hOc4la, hOc4v, FG1, and FG2) and ventral (conjunction of bilateral FG3 and FG4) components of OTC (Eickhoff et al. 2005). Within the two masked areas we used a threshold of FWE = .05, and then lowered the threshold to uncorrected p = .001, if no activity was detected, or if it was detected in only one hemisphere. This procedure resulted in six functionally defined ROIs: LOTC-body, LOTC-face, LOTC-object, VOTC-body, VOTC-face, and VOTC-object. To define EVC, we used the posterior (i.e. most foveal) ~2/5 of the V1 mask as defined by the Anatomy Toolbox.

### Representational similarity analysis

RSA was used to compare the activity patterns from the different ROIs to the stimulus models, similarity judgments and the layers of a suite of DNNs (Kriegeskorte and Kievet, 2013; Kriegeskorte, Mur, and Bandettini, 2008). For each comparison RDMs were constructed, which are matrices that are symmetrical around the diagonal and reflect the pairwise dissimilarities between all stimulus conditions. Matrices from different data modalities can be directly compared in order to evaluate the second order isomorphisms of the dissimilarities between conditions. RSA was carried out using CoSMoMVPA, along with custom code (Oosterhof et al 2016).

Neural RDMs for the different ROIs, for each subject, were constructed using the (non-cross-validated) Mahalanobis distance as the dissimilarity metric, characterized as the pairwise distance along the discriminant between conditions for the beta weight patterns in an ROI (Ritchie and Op de Beeck, 2019; Walther et al. 2016). To assess the between-subject reliability of these RDMs, the RDM of one subject was left-out, and those of the remaining subjects averaged and Pearson’s *r* correlated with the left-out subject’s RDM. This was carried out for all subjects, and the resulting coefficients averaged. The resulting average value was used as an estimate of the noise ceiling when correlating individual neural RDMs with the RDMs from the other data modalities. Visualization of group averaged neural RDMs included multi-dimensional scaling (MDS) with stress 1 as the criterion.

For the main model RDMs, a 54-value vector was coded based on whether a stimulus was animate or not (Animacy), a face-body or not (Face-Body), or its rank (Mammal 1 = 1; invertebrates = 6) in the intuitive taxonomic hierarchy (Taxonomy). The values of the model RDMs were then filled based on the absolute difference in the pairwise values in these coding vectors. To construct an RDM from GIST, each image was segmented into a 4 × 4 grid, and Gabor filters (8 orientations and 4 spatial frequencies) were applied to each block in the grid. For each image, the values for each filter were converted to a vector, and all pairwise 1 – *r* Pearson’s correlations between these vectors were used to fill the cells of the RDM. For the different DNNs (described below), layer-specific RDMs were constructed based on the 1 – *r* pairwise Pearson’s correlation between the vectors of unit responses for each image. For the similarity judgments with a 6-point scale, RDMs were constructed based on the absolute difference in judgments along the scale. For the other similarity tasks participants’ responses resulted in a ranking vector (averaged across runs in the case of the in-scanner preference task), and the difference in rank was used as the dissimilarity metric to construct the matrices. Individual behavioral RDMs were averaged to create a group-averaged matrix for comparison with individual neural RDMs. To assess the reliability of the group-averaged human similarity RDMs, the individual RDMs were split into two groups, then averaged and correlated. This was done for all possible half-splits of the data, and the resulting average coefficient value was transformed using the Spearman-Brown formula. This resulting value gives an estimate of the reliability of the group-averaged data, based on the full sample size (DiCarlo and Johnson 1999; Op de Beeck et al., 2008).

To compare RDMs from different data modalities, the bottom half of each matrix was converted to a vector and the Spearman rank-order correlation was calculated between matrices. The median correlations across subjects were tested for significance using the Wilcoxon Signed Rank Test. Since the test statistic W can be computed exactly for N ≤ 15, it is reported along with the p-value. In the case of the DNNs, in which several comparisons of the same type were made, the false discovery rate (FDR) adjusted p-values are reported to correct for multiple comparisons.

### Commonality analysis

Commonality analysis is a method for determining whether predictors uniquely or jointly explain variance in the dependent variable (Newton and Spurell, 1967; Siebold, McPhee, 1979). This method has increasingly been used in conjunction with RSA and other MVPA methods and is also sometimes known as “variance partitioning” (Groen et al. 2018; Hebart et al. 2018; Lescroart et al. 2015). In the present case of three predictors (a, b, c) for some dependent variable y, there will be seven coefficients of determination (*R*^2^) for all possible combinations of predictors in a linear regression model: *R*^2^_y ∙ a_, *R*^2^_y ∙ b_, *R*^2^_y ∙ c_, *R*^2^_y ∙ ab_, *R*^2^_y ∙ ac_, *R*^2^_y ∙ bc_, *R*^2^_y ∙ abc_. The last of these is the “full” model, for which the variance is partitioned based on differential weighting of the *R*^2^ of the different models. When there are only three predictors, the partitioning can be performed using a simple weighting table (Nimon and Reio, 2011, Table 2), in which the vector of coefficients is multiplied with row vectors of weights for each of the unique and common variance components. These results were visualized with EulerAPE (Micallef and Rodgers, 2014), which can be used to plot overlapping ellipses proportional to the variance partition of the total explained variance (Groen et al. 2018). While in principle negative variance can reflect informative relationships between predictors (Capraro and Capraro, 2001), in the present context these values were typically so small (e.g. −0.1 % of the explained variance) as to be negligible. They were therefore excluded from the visualization. Notably, the exclusion of these negative values entails that the displayed values in the Euler plots depicting the commonality analyses that were performed will not sum to exactly 100 %, as is normally the case when all unique and common variance components are combined (Nimon and Reio, 2011).

To carry out the multiple regression necessary for commonality analysis RDMs were converted to vectors, as with RSA, and the group-averaged neural dissimilarity values were regressed on the different model or behavioral dissimilarity vectors. Since this application of linear regression violates standard assumptions of independence between samples and normality, significance for the full model was determined by using a permutation test, in which the values of the group-averaged neural dissimilarities were shuffled 1000 times, and the full model was fit to the shuffled vectors. The resulting proportion of *R*^2^–values greater than that observed for full model (fit to the un-shuffled dissimilarities) provides the p-value for the test.

### Univariate analysis

For each subject the beta weight values were averaged across all runs, and then all voxels, for each stimulus condition. Univariate RDMs were then constructed based on the pairwise absolute differences in the beta weight values for each condition. The individual subject univariate RDMs were then correlates with the main model matrices as with the other forms of RSA described above. To further visualize the univariate results, average beta weight values were calculated for each of the clusters of four face and body images for each level of the Taxonomy model, and for all 6 of the natural object images.

### Deep convolutional neural networks (DNNs)

Networks consisted of stacked multiple convolutional layers (conv) that were intermittently followed by pooling operations, which fed into FC layers prior to output. Each DNN was pre-trained on the ImageNet dataset (Russakovsky et al. 2015). To generate the response vectors for RSA we passed each image through the networks, with the activation weights of each layer as outputs. Softmax classification layers, which reflect the 1000 labels for image net, were excluded from the analysis. The networks used in our analysis are well-known for their performance in the ImageNet competition, or in the case of CORnet, has been promoted as superior in brain predictability, in contrast to image classification performance, using Brain-Score (Schimpf et al. 2018).

Caffenet is an implementation of the AlexNet architecture within the Caffe deep learning framework (Krizhevsky, Sutskever, and Hinton, 2012; Jia et al. 2014). The network includes 5 conv layers followed by 3 FC layers, for a total network depth of 8 layers. VGG-16 consists of 13 conv layers interspersed with 5 pooling layers, followed by 3 FC layers (Simonyan and Zisserman, 2015). GoogLeNet or InceptionNet is a 22-layer deep network, when counting only the parameterized layers (Szegerdy et al., 2015). As with the majority of standard networks, the initial layers are conventional conv layers followed by max pooling operations, and the final layers involve (average) pooling followed by a single FC layer. The main architectural point of difference from standard serial networks is that in GoogLeNet, intermediary layers consist of stacked “inception” modules, which are themselves miniature networks containing parallelized conv and max pooling layers, with convolutions of different sizes. ResNet50 is a deeper network (50 layers deep), with 48 convolutional layers and 2 pooling layers (He et al. 2015). The unique feature of ResNet50 is the implementation of shortcut connections that perform identity (residual) mappings between every 3 layers. CORnet is a family of architectures that include recurrent and skip connections, with all networks containing 4 layers that are pre-mapped onto the areas of the ventral visual pathway in the primate brain: V1, V2, V4, and IT (Kubilius et al. 2018). We used CORnet-S, which combines skip connections with within-area recurrent connections to perform best “overall” on Brain-Score (Schimpf et al. 2018), and so in principle should potentially be superior at matching the dissimilarity structure of OTC.

## Acknowledgements

J.B.R. received funding from the FWO (Fonds Wetenschappelijk Onderzoek) and European Union’s Horizon 2020 research and innovation programme under the Marie Skłodowska-Curie grant agreement No. 665501, via an FWO [PEGASUS]^2^ Marie Skłodowska-Curie fellowship (12T9217N). A.A.Z. and H.P.O. were funded by grant C14/16/031 of the KU Leuven Research Council. Neuroimaging was funded by the Flemish Government Hercules Grant ZW11_10.

## Author Contributions

Conceptualization: J.B.R. and H.P.O.; Methodology: J.B.R. and H.P.O.; Investigation: J.B.R, A.A.Z, J.B., S.S., and K.V.; Writing – Original Draft: J.B.R and H.P.O; Writing – Reviewing & Editing: J.B.R, A.A.Z., and H.P.O; Funding Acquisition: H.P.O; Resources: H.P.O.; Supervision: J.B.R. and H.P.O.

---

**Figure S1.**
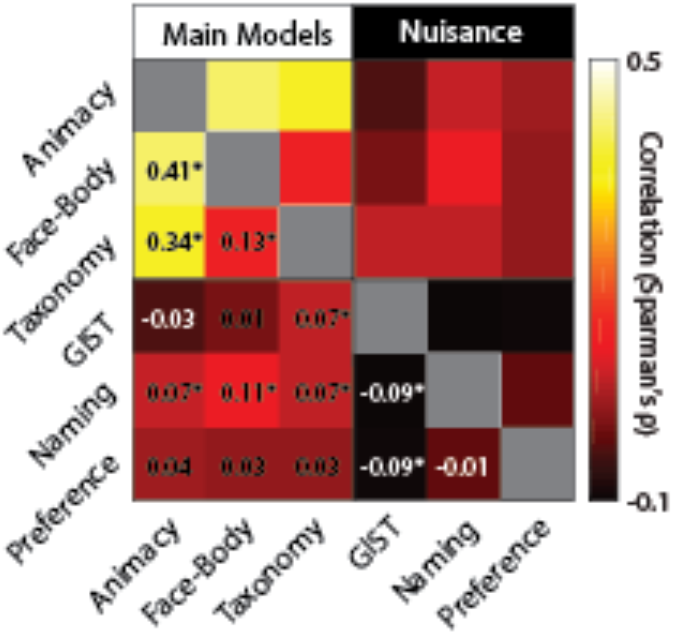
Correlation between main model and nuisance predictor RDMs. Cells in the matrix correspond to the pairwise correlations (Spearman’s ρ) between matrices. Text for negative correlations appear in white. * = p < 0.05. Note: if the natural object images are excluded, the Animacy RDM contains no internal structure, and the Face-Body and Taxonomy RDMs are not correlated (ρ = −0.03, p = 0.28).

**Figure S2.**
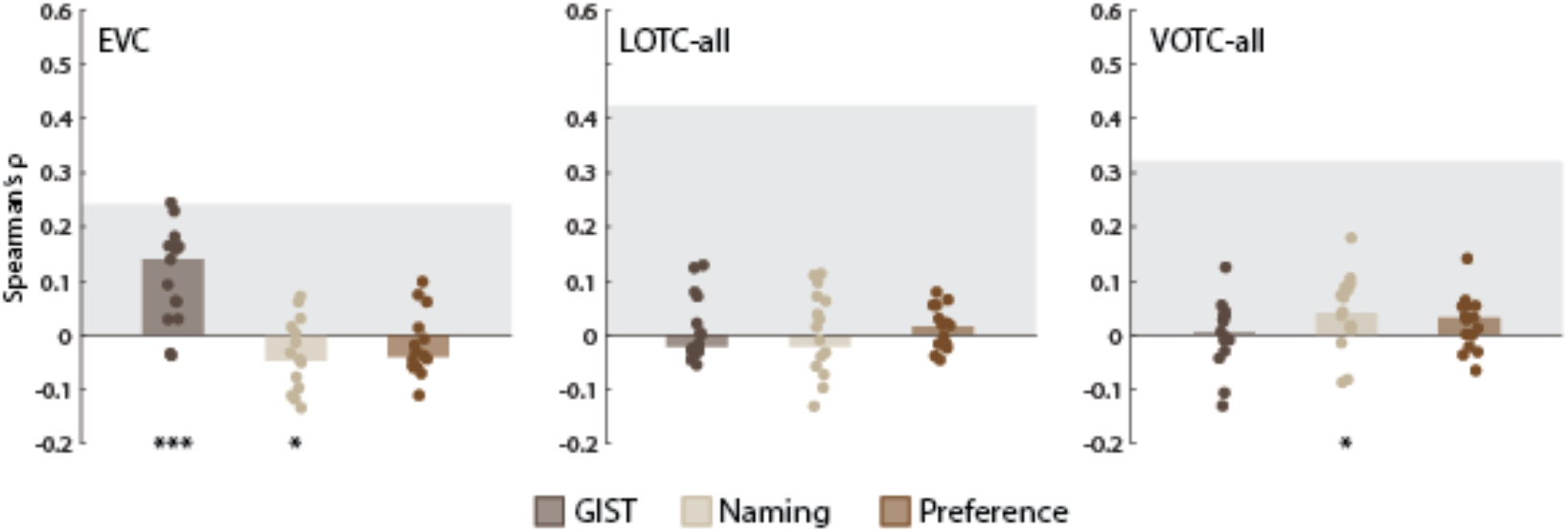
Comparing the nuisance model RDMs to EVC, LOTC-all, and VOTC-all. **(A)** Bar charts indicating the median Spearman’s ρ rank-order correlations between neural RDMs for individual subjects (dots) and the model RDMs. Gray blocks indicate the noise ceiling. * = p < 0.05 based on Wilcoxon Signed Rank Tests. As the naming RDM was correlated with those for the Face-Body and Taxonomy models, we carried out partial correlations of these RDMs with the individual neural RDMs for LOTC-all and VOTC-all, controlling for the naming RDM. In both cases, the median correlations decreased slightly after controlling for the naming RDM (both: Δ median ρ < 0.01), and remained highly significant (both: W(15) = 120, p = 6.10e-05).

**Figure S3.**
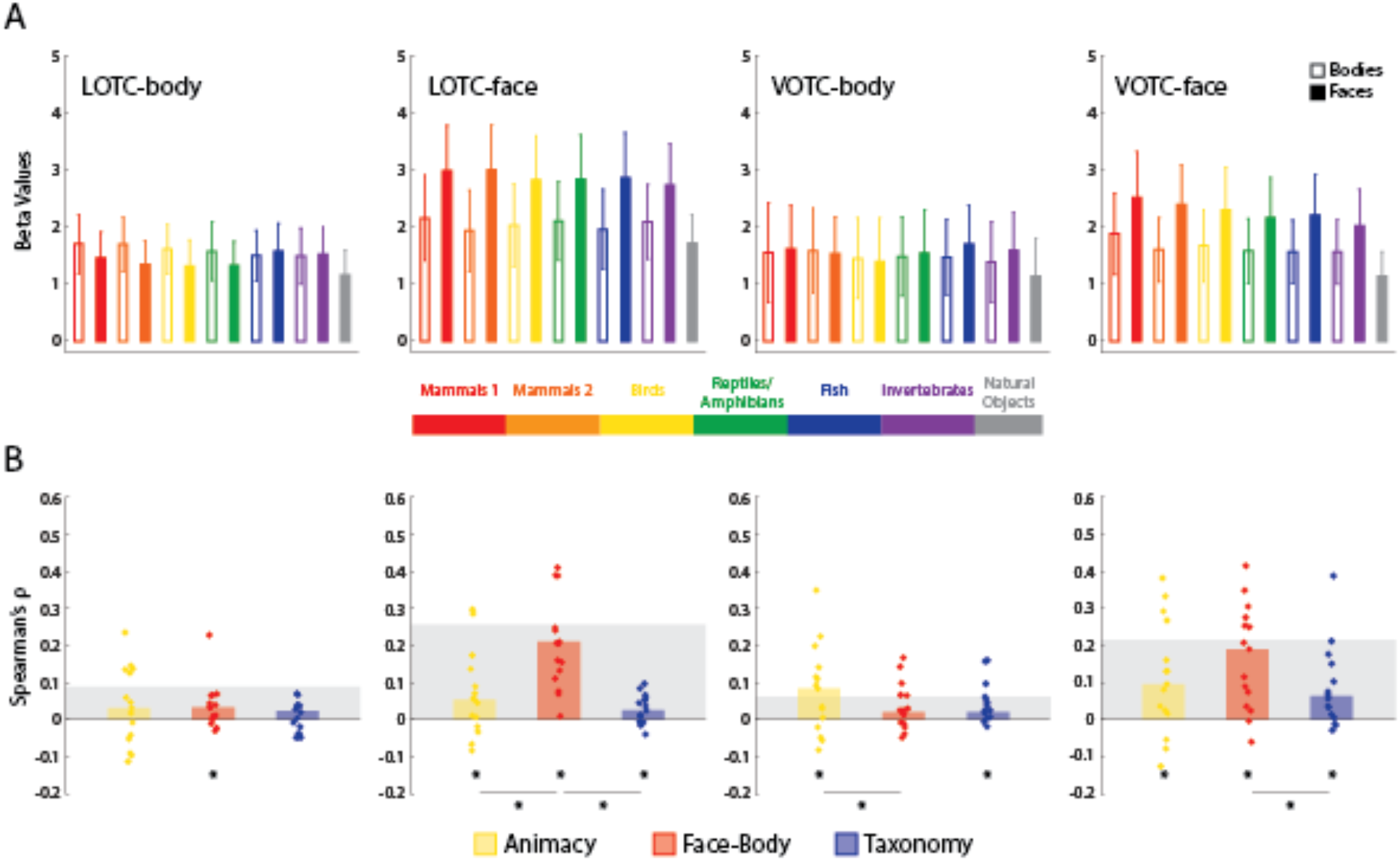
Results of univariate RSA for the face- and body-selective ROIs. **(A)** The average univariate responses for face and body images, at each level of the taxonomic hierarchy, are depicted against the average data of natural object images. Error bars indicate the standard error of the mean (SEM). **(B)** Bar charts indicating the median Spearman’s ρ rank-order correlations between univariate RDMs for individual subjects (dots) and the three main model RDMs, for all four ROIs. Gray blocks indicate the noise ceiling. * = p < 0.05 based on Wilcoxon Signed-Rank Tests.

**Figure S4.**
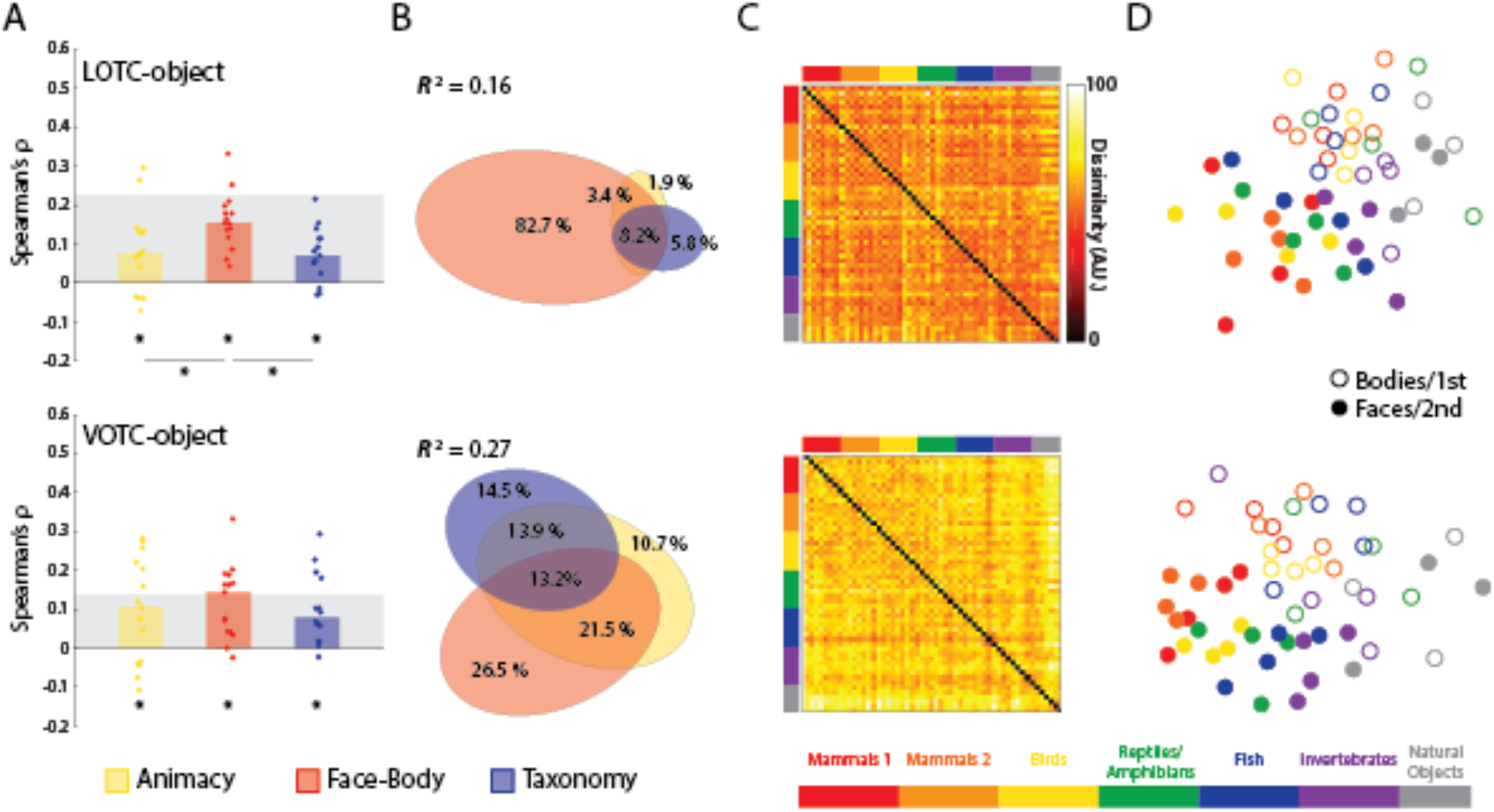
Results of model RDM comparisons to object-selective regions of OTC. **(A)** Bar charts indicating the median Spearman’s ρ rank-order correlations between neural RDMs for individual subjects (dots) and the three main model RDMs, for all four ROIs. * = p < 0.05 based on Wilcoxon Signed Rank tests. **(B)** Results of commonality analysis for all ROIs, visualized with Euler diagrams. Coefficients of determination (*R*^2^) are for the full regression models and indicate the total percentage of explained variance. **(C)** Group-averaged neural RDMs for the four ROIs, with the axes color-coded based on the taxonomic hierarchy. Dissimilarity values are scaled to range between 0 and 100. **(D)** 2D multi-dimensional scaling applied to the dissimilarity matrices for each ROI. Points are color-coded to reflect the taxonomic hierarchy and are either rings or dots to reflect the face/body division, or the 1^st^/2^nd^ item for each natural object type.

**Figure S5.**
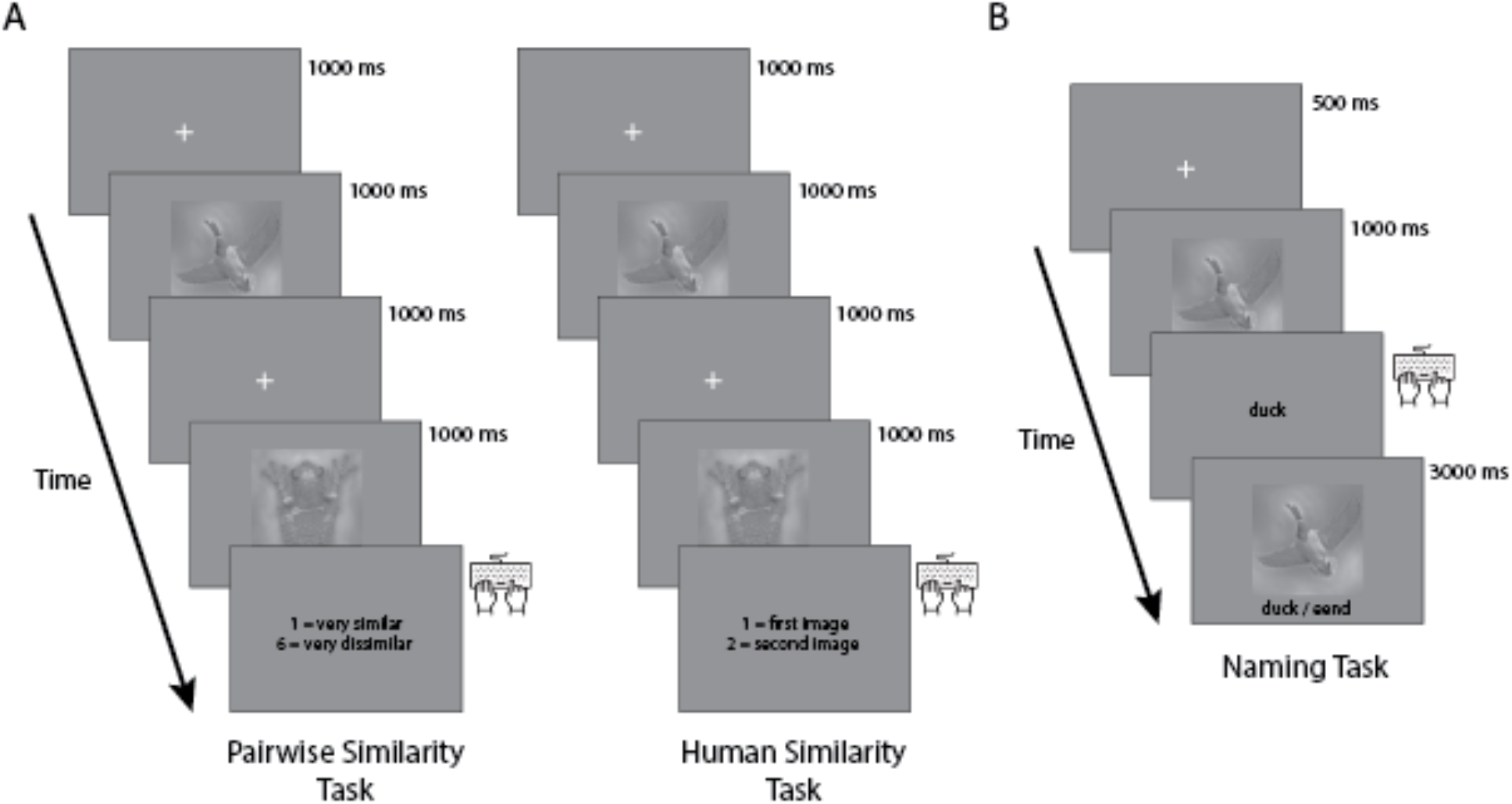
Structure of similarity judgment and naming tasks. **(A)** Single trial of the pairwise (face/body) similarity and human (face/body) similarity tasks. Depiction is for a single trial for the body images, for each type of similarity judgment. **(B**) Single trial of the naming task. Depiction is for a single trial for one of the animal images.

